# Associations and recovery dynamics of the nasopharyngeal microbiota during influenza-like illness in the aging population

**DOI:** 10.1101/2021.09.14.458655

**Authors:** Sudarshan A. Shetty, Josine van Beek, Elske Bijvank, James Groot, Sjoerd Kuiling, Thijs Bosch, Debbie van Baarle, Susana Fuentes

**Author notes:** **Corresponding author:** Susana Fuentes.

## Abstract

**Background:** Older adults are more susceptible to respiratory pathogens, several of which have been associated with an altered respiratory microbiota. Influenza-like illness (ILI), a disease caused by respiratory pathogens including but not exclusively by influenza virus, is a major health concern in this population. However, there is little information on changes in the nasopharyngeal (NP) microbiota of older adults associated with respiratory infections identified by/ reported as ILI, as well as its dynamics during recovery. Here, we compared the NP microbiota in older adults who presented with ILI (n= 240) to the NP microbiota in older adults not reporting an ILI event (n= 157) during the 2014-2015 influenza season. To investigate the dynamics of the microbiota from the acute phase to the recovery phase of the infection, participants reporting an ILI event were sampled at onset of infection (<72 hours), at 14 days and at 7-9 weeks after infection (recovery sample).

**Results:** Cross-sectional analysis of the microbiota at the different time-points showed no differences in alpha diversity between the groups. A small but significant effect of the ILI was observed on the microbiota community and structure when compared to controls and recovery samples. Furthermore, the NP microbiota exhibited inter-individual differences in dynamics from onset of ILI to recovery. *Corynebacterium*, one of the keystone species in the upper respiratory tract, was negatively associated with ILI and its abundance increased after recovery. Potential pathobionts such as *Haemophilus, Porphyromonas* and *Gemella* had higher abundances during acute-ILI. Stability and changes in the NP microbial community showed individual dynamics. Key core genera, *Corynebacterium, Moraxella* and *Dolosigranulum* exhibited higher inter-individual variability in acute-ILI, but showed comparable variability to controls after recovery. Participants in the ILI group with higher core microbiota abundances at the acute phase showed higher microbiota stability after recovery.

**Conclusions:** Our findings demonstrate that acute-ILI is associated with small but significant alterations in the phylogenetic structure of the NP microbiota in older adults. The observed variation in the core microbiota suggests potential imbalances in the ecosystem, which could play a role in the recovery of the NP microbiota after an ILI event.

## Background

Respiratory viral infections are a major cause for public health concern, especially due to their mode of transmission and high risk of morbidity and mortality in at risk populations such as older adults [1]. While vaccinations have the potential to prevent infections, their efficacy can vary based on several factors, such as ageing and the subsequent deterioration of the immune system [2-6]. Influenza-like illness (ILI) refers to a defined set of symptoms associated to the disease caused by respiratory bacterial or viral pathogens, including but not exclusively by influenza virus. [7].It is crucial to understand the factors that may play either a protective or pathogenic role in this respiratory disease, such as that of the commensal respiratory microbiome.

The respiratory tract harbors between 10^2^ to 10^6^ bacteria depending on the site, with the nasopharynx (NP) consisting of ∼10^3^ of bacteria [8]. The upper respiratory tract microbiota of older adults differs from that of middle-aged healthy adults [9]. These differences have been associated with, among others traits of ageing, immune-senescence, i.e., a dysregulation of the immune system occurring with age suggesting a potential role of the microbiota through interaction with the local immune response [9-12]. Commensal NP microbiota, can play an important role in inhibiting colonization and expansion of invading pathogens by exhibiting colonization resistance similar to what occurs in other host-associated niches, [13]. E.g. for bacterial infections, colonization of *Staphylococcus lugdunensis* (a lugdunin producing bacterium) was shown to reduce *Staphylococcus aureus* carriage and could play a role in preventing staphylococcal infections [14]. Studies investigating the upper respiratory tract microbiota during influenza virus infection have identified differences between infected and non-infected individuals [15, 16]. However, to our knowledge, information on changes in the NP microbiota during the onset of ILI and its dynamics during recovery after an ILI event in the ageing population remains under explored. A better understanding of microbiota dynamics may provide clues for potential microbial markers of recovery from ILI and susceptibility to secondary bacterial infections such as pneumonia, which is of special relevance in the ageing population.

We investigated the NP microbiota in older adults reporting an ILI event in comparison to NP microbiota in participants not reporting an ILI event throughout the 2014-2015 influenza season, considered as controls [6]. We used 16S rRNA gene sequencing to assess the composition of the NP microbiota in a subset of participants (n= 397) from the ILI cohort [6]. Controls were sampled throughout the season equally distributed over the different age groups at two timepoints, 14 days apart. Participants reporting an ILI event were sampled three times: at onset, at 14 days and at 7-9 weeks, considered as the recovery sampling phase. We observed differences in NP microbiota community composition between the ILI and control groups, indicating changes associated with acute infection. In longitudinal samples from ILI participants, the NP microbiota during recovery differed markedly from that observed during the acute phase. Our study highlights the need for further mechanistic and longitudinal studies to understand the role of the NP microbiota and its link with susceptibility as well as recovery from ILI in the ageing population.

## Material and Methods

### Study design and participants

The participants (n= 397) in this study were part of the prospective surveillance cohort of individuals with influenza-like illness (ILI) which was previously described [6, 17]. For this study, we included all individuals that had nasophyrangeal swabs available and reported ILI symptoms once during the 2014-2015 season as well as a control group consisting of participants with no ILI reported in the same season (see supplementary fig. 1). Detailed information of participants in this study is provided in Table 1. The differences in sex, age and body-mass index were tested using Fisher test, t-test, sum statistic (without blocking by stratum i.e., age and gender). For characteristics such as medications, comorbidities, presence of pathogens in the cohort we used Cochran-Mantel-Haenszel test with blocking per strata, where stratum is defined as the interaction of sex and age groups (5-year range between 60-90 years). Ethical clearance was obtained from the Medical Ethical Committee Noord Holland and written informed consent was provided by the participants. This study is registered with the Netherlands Trial Registry, number NL4666.

**Table 1:** Baseline information from participants in this study. Asterisks indicate differences detected between the control and ILI groups: ****FDR < 0.0001, ***FDR < 0.001, **FDR <0.01; *FDR<0.1.

|  | Control | ILI | Statistical significance |
| --- | --- | --- | --- |
| Total participants | N = 157 | N = 240 |  |
| Participants with longitudinal samples | N<sub>(baseline + 14days)</sub> = 78 | N<sub>(acute + 14 day + recovery)</sub> = 81 |  |
| Age | 71.4 (± 6.2) | 69.5 (± 6.0) | 0.004 (T test) |
| BMI | 26.5 (± 4.4) | 25.4 (± 3.7) | 0.009 (Sum-statistic) |
| Gender (M/F) | 80/77 | 115/125 |  |
| Antibiotics in 2014 (Yes/No) | 16/141 | 68/172 | **** |
| Antimicrobials in 2014 (Yes/No) | 19/137 | 68/172 | **** |
| Co-morbidities 2014~ (Yes/No) | 58/99 | 107/133 | * |
| Autoimmune disease | 4/153 | 13/227 |  |
| Chronic cardiovascular disease | 22/135 | 38/202 |  |
| Diabetes | 17/140 | 21/219 |  |
| Malignancies | 15/142 | 10/230 | * |
| Respiratory disease | 17/140 | 53/187 | ** |
| Smoking | Active=7 | 19 |  |
|  | Passive=7 | 5 |  |
|  | Non=143 | 216 |  |
| Influenza vaccination (2014-15) | Yes = 123 | 161 | * |
|  | No = 32 | 78 |  |
|  | Unknown= 2 | 1 |  |
| Medication |  |  |  |
| ACE inhibitors | 21/136 | 25/215 |  |
| Alpha-/betablocker | 52/105 | 44/196 | ** |
| Analgesic | 27/130 | 62/178 | * |
| Anticoagulants | 36/121 | 43/197 |  |
| Antidepressants | 16/141 | 8/232 | ** |
| Antiepileptic | 3/154 | 7/233 |  |
| Antihistamines | 5/152 | 10/230 |  |
| ARBs | 29/128 | 32/208 |  |
| Asthma/COPD | 9/148 | 26/214 | * |
| Calcium medication | 22/135 | 29/211 |  |
| Corticosteroids | 24/133 | 56/184 | * |
| Benzodiazepine | 10/147 | 5/235 | * |
| Insulin | 2/155 | 6/234 |  |
| PPIs | 49/108 | 59/181 |  |
| Statins | 56/101 | 54/186 | ** |

~Participant has a comorbidity (respiratory, cardiac, diabetes, kidney, transplant, autoimmune, asplenia, leukemia, lymphatic cancer, malignancy)

### Sample collection

Nasopharyngeal (NP) swabs were collected between Oct 1, 2014, and June 15, 2015, during the 2014–15 influenza season from participants registering an ILI event within 72 h of reporting symptoms (acute-ILI; visit-1), after 14 days (ILI-14days; visit-2) and 7–9 weeks after the ILI event (recovery sample; visit-3). The Dutch Pel criteria was used to determine Influenza-like Illness, where having a fever (≥37·8°C) with at least one other symptom such as headache, myalgia, sore throat, coughing, rhinitis, or chest pain was used to classify individuals into the ILI group [6]. Causative agents were identified using NP and oropharyngeal (OP) swabs. NP swabs were further used for microbiota profiling. Participants without any reported ILI event were used as controls and randomly sampled throughout the season (visit-1) and 14 days after their first visit (visit-2).

### DNA extraction and sequencing

The DNA extraction protocol for low-biomass samples that was previously demonstrated to be robust was used to extract DNA from NP swabs with slight modifications [18]. We used the modified Agowa Mag DNA extraction kit (LGC genomics, Berlin, Germany). For each batch of DNA extraction, the ZymoBIOMICS Microbial Community Standard (Zymo catalog number: D6300) was 1000x diluted and 200 µl of this diluted Zymo mock was included as positive control together with two negative controls containing only the lysis buffer. The NP FLOQSwabs® in Copan’s Liquid Amies Elution were thawed on ice and 200 µl vortexed for 10 seconds. For each sample, 600 µl of lysis buffer containing zirconium beads (diameter 0,1 mm, Biospec Products, Bartlesville, OK, USA) and 550 µl phenol (VWR International, Amsterdam, the Netherlands) were added. The bead-beating step was done twice for 2 minutes at 3,500 oscillations/minute by bead beating (Mini-Beadbeater-24, Biospec Products). Between each bead-beating step, the tubes were transferred on ice for 2 minutes. The tubes were then centrifuged for 10 minutes at 4,500 x g and the clear aqueous phase was pipetted and transferred to a new 2 ml Eppendorf tube that consisted of 1.3 ml binding buffer and 10 µl magnetic beads. The 2 ml Eppendorf tube was kept for shaking for 30 minutes at 900 RPM and then placed on a magnetic separation rack. The clear liquid was discarded and the magnetic beads were washed and dried for 15 minutes at 55°C. The DNA was eluted in 35 µl elution buffer.

The rRNA V4 region of the 16S rRNA gene was amplified by PCR using the 515F (5’-GTG CCA GCM GCC GCG GTA A-3’) and 806R (5’-GGA CTA CHV GGG TWT CTA AT-3’) primers including the Illumina adapters and sample specific barcodes [19, 20]. For PCR, we included additional DNA mocks, ATCC-MSA-2004 and an in-house DNA mock consisting of *Haemophilus influenzae, Streptococcus pneumoniae, Streptococcus pyogenes* (group B), *Klebsiella oxytoca, Klebsiella pneumoniae*, Hemolytic *Streptococcus* Group A, *Pseudomonas aeruginosa, Staphylococcus epidermidis, Staphylococcus aureus* and *Moraxella catarrhalis* pooled in equal ratios based on 16S rRNA gene qPCR data. Overall, DNA blanks, non-template controls (NTC), ZymoBIOMICS Microbial Community Standard (whole cell) and gDNA mocks were included in each PCR plate and sequenced alongside the samples sequenced with an Illumina MiSeq instrument (Illumina Inc., San Diego, CA, US) following manufacturer’s recommendations.

### Identification of amplicon sequence variants (ASVs)

All the bioinformatic analysis were done in R (v3.6.0) and RStudio (v1.1.383), unless otherwise stated [21, 22]. For each sequencing run, raw reads were filtered, trimmed and denoised into amplicon sequence variants (ASVs) using the dada2 R package (v1.14.1) [23] using default parameters, except for *filterAndTrim* for which we used following parameters (truncLen=c(200,150), trimLeft = c(20,22)). The creation of the ASV table was followed by removal of chimera and ASV tables were merged from individual sequencing runs.

The ASVs were assigned taxonomy using the RDP classifier and SILVA database v138 with *assignTaxonomy* (key parameters: minBoot=80 and tryRC=TRUE). Species level assignments were done using the *addSpecies* function [24, 25]. The resulting ASV table and taxonomic table were combined with sample data into a phyloseq (v1.30) object for downstream analysis [26].

### Removal of potential contaminant ASVs

In order to identify and deal with contamination from exogenous sources such as reagents, of special relevance in our low biomass samples, we included several positive and negative technical controls. The technical controls were as follows: 1) DNA extraction controls (n = 32) 2) PCR non-template controls (n = 13) and three different mock communities. Genomic DNA mixtures included an ATCC mock community (n = 15), and in-house mock community (n = 10) and the ZymoBiomics mock community (n = 53) including genomic DNA as well as whole-cell mixtures. As a first step to remove contaminant ASVs we used the prevalence methods (threshold 0.5) from the decontam R package function *isNotContaminant()* [27]. In contrast to the widely used, *isContaminant* approach, the *isNotContaminant* is stricter and focuses on identifying taxa that are not likely to be contaminants [27]. ASVs over-abundant in negative controls compared to true samples, and those classified as Cyanobacteria, Chloroflexi and Mitochondria were removed. Next, a co-abundance approach was used, by applying a correlation-based clustering approach for the identified blank-sample ASVs in all the data (technical controls plus NP samples). Optimal number of clusters of co-occurring taxa were identified using gap statistic with 500 permutations with *fviz_nbclust()* function in factoextra R package (v1.0.7) using the *hcut* method and visualized using *aheatmap()* function in the NMF R package (v0.22.0) [28, 29]. To check for consistency in the 9 clusters that were identified, we split the data into training and test data by randomly subsampling 500 samples and testing clustering with mantel test function in vegan R package (v2.5.6) [30].

Finally, to reduce extremely rare ASVs, we aggregated data at the phylum level and excluded the following phyla based on low abundance (< 0.02) and prevalence (<25%) in all samples combined: Synergistota, Planctomycetota, Acidobacteriota, Spirochaetota, Bdellovibrionota, Verrucomicrobiota, Gemmatimonadota, Desulfobacterota, Myxococcota Crenarchaeota, Abditibacteriota, Euryarchaeota, Halobacterota, Armatimonadota, Fibrobacterota, Nitrospirota, Dependentiae. The Bacterial Diversity Metadatabase (BacDive; https://bacdive.dsmz.de/advsearch) was used to search for Genus and species epithet for the most dominant/prevalent ASVs from Fusobacteriota and Campilobacterota to confirm the source of isolation of nearest cultured representative [31]. Despite being prevalent (63%) and constituting 0.15% of the reads, we removed (as a rational choice) phylum Deinococcota, because ASVs from this phylum belonged to the *Deinococcus-Thermus* group, identified as reagent contaminants in several studies [32-35].

### Microbial community analysis

Phylogenetic diversity, Simpson’s evenness and beta-diversity (Generalized UniFrac and unweighted Unifrac) analyses were performed on rarefied data subsampled at 1932 reads/sample (86.5% of ASVs were retained), using R packages picante (v1.8.1), microbiome (v2.1.1) and MiSPU (v1.0) [36-40]. Association between beta-diversity and health status was assessed using a non-parametric Analysis of similarities (ANOSIM, vegan R package) with 999 permutations. Within group differences in beta-diversity (divergence) and intra-individual stability (1-GUniFrac) in microbiota was calculated based on the GUniFrac. Proportional variability (PV) in core genus relative abundances was calculated as described previously [41, 42]. To avoid instances where values are divided by zero, due to either detection limit or true absence of taxa in samples, we add a small constant value (1% of mean relative abundance) to relative abundances of each taxon before calculating PV. To avoid biases resulting from differences in number of samples between groups, we sampled 80% of the samples in each group with 999 bootstrap iterations. Taxonomic compositions were visualized using microbiomeutilities (v1.00.12), pheatmap (v1.0.12) and ggplot2 (v3.3.3) R packages [43, 44]. Between group alpha diversity, divergence and intra-individual stability were compared using the Wilcoxon tests within the *stat_compare_means* function in the R package ggpubr (v0.2.5) [45].

### Associations with microbiota

Associations between microbiota composition and variables such as ILI status, pathogen type (bacterial or viral), medications and co-morbidities were carried out using the procedure described previously [17]. Briefly, ASVs were agglomerated to genus level and those genera with minimum 0.001 relative abundance in 10% of the samples were selected for testing associations mentioned above. The sum-statistic was used for testing associations between genera relative abundances and numerically coded variables [46]. The R package coin (v1.3-1) was used for statistical testing [47]. To reduce the effect of confounding variables such as participant age and gender, these were stratified as blocks. The Benjamin-Hochberg procedure was applied for multiple testing to each sub-study [48].

## Results

### Cohort description

An overview of the study design and analysis is given in supplementary fig. 1. 397 participants, with a mean chronological age of 70.3 (±6.2) years were selected for this study based on availability of nasopharyngeal swabs. Of these, 157 participants (approx. 40%) did not report an ILI-event throughout the 2014-2015 influenza season, and were therefore used as controls, and 240 reported an ILI event, and were considered as the ILI group. An overview of participant information is provided in Table 1. While the majority of participants in the control (89.2%) and ILI (77.9%) groups had no additional respiratory diseases (such as asthma or COPD) at the time of sampling for this study, these were significantly higher in the ILI group.

Participants from the control group had higher rates of influenza vaccination (78.3%) than the ILI group (67.1%). Comorbidities were more frequent in the ILI group, as well as use of antibiotics and analgesics (Table 1). The most common causative agents for ILI were *Influenza* virus (21.2%), *Rhinovirus* (17.5%), *Haemophilus* (15.4%), and *coronavirus* (7.9%) (Supplementary Table 1).

### Microbial community composition in the NP in the ageing population

To investigate the composition of the NP microbial community of the study participants, we first removed contaminant sequences due to high prevalence of taxa previously reported as reagent contaminants, e.g., *Ralstonia, Bradyrhizobium, Mesorhizobium, Comamonadaceae* (Supplementary fig. 2). After removing these contaminants, the top three most abundant phyla, namely Actinobacteriota, Firmicutes, and Proteobacteria, accounted for 97.8 % of the total composition in all samples (Supplementary Table 3). The top five genera, *Corynebacterium, Moraxella, Dolosigranulum, Staphylococcus* and *Haemophilus* contributed to 83.8% of the total (Supplementary Table 4). ASVs unclassified at the genus level contributed to 3.11% of the total. Based on genus abundance across samples, C*orynebacterium* was dominant in 48.9% of the samples, *Moraxella* in 20.5%, *Staphylococcus* in 13.9%, *Dolosigranulum* in 5.2%, and *Haemophilus* in 4% of the samples (Supplementary Table 5). In 47% of the samples, a single ASV was contributing to more than 50% of the total abundance. This suggests that the majority of ASVs detected in an individual’s NP microbiota were of low abundance.

### Effect of ILI on the NP microbiota diversity and structure

To assess differences in microbial diversity and community evenness, we compared the NP microbiota between controls and the ILI group throughout the different phases of the ILI. Comparison of phylogenetic diversity (Faith’s Phylogenetic diversity, PD) revealed higher diversity in samples 14 days after ILI when compared to controls, acute-ILI and samples after recovery (Fig. 1A). We observed that the NP microbiota exhibited low evenness (Simpson’s evenness, Fig.1A), supporting our previous observation of single ASVs dominance in the NP microbiota, with higher levels observed in controls when compared to samples from 14-days after ILI. However, significance of these differences disappeared after correcting for multiple testing.

**Figure 1:**
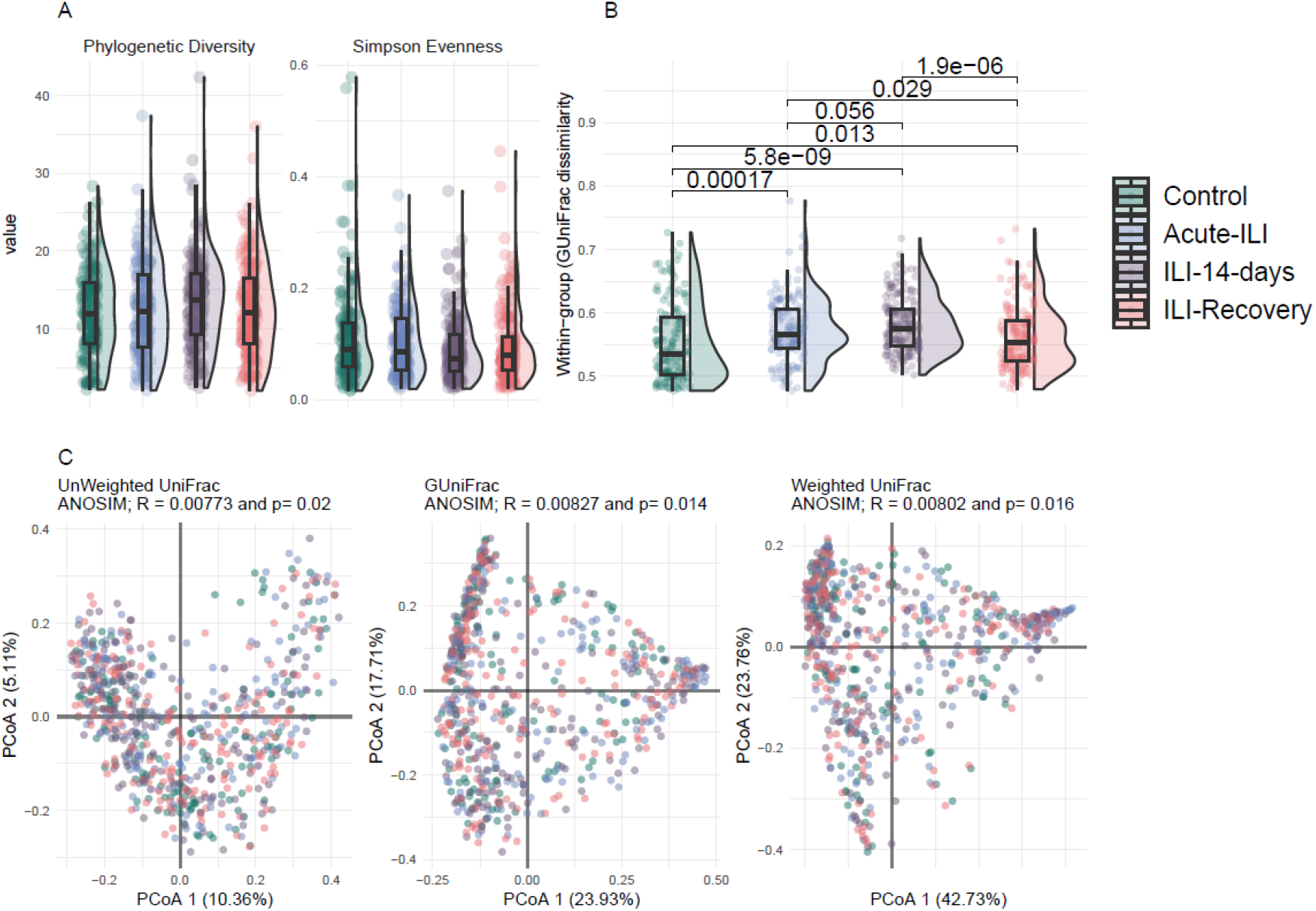
Comparison of diversity between groups. A] Alpha diversity measures, phylogenetic diversity and evenness comparisons between groups. B] Inter-individual variation in community composition based on generalized UniFrac distances between samples within a group. Statistical comparisons are based on Wilcoxon rank-sum test, corrected for multiple comparisons using Benjamini-Hochberg Procedure. C] Comparison of beta-diversity between groups based on Unweighted, Generalized and Weighted UniFrac distances.

Within-group variation in the overall microbiota community composition and structure was different between controls and the ILI group at all time-points (Fig.1B). This variation was higher in samples from the acute-ILI and at 14 days after when compared to controls and samples from the recovery phase. Due to the observed unevenness in the NP microbiota (Fig.1A), we used three distance measures, i.e. unweighted, generalized and weighted UniFrac, to account for mono-dominance and rarity in NP microbiota. The microbiota composition and structure were significantly different between the control and ILI groups (Fig. 1C, Unweighted UniFrac, ANOSIM *R* = 0.007, *P*_anosim_ = 0.02; GUniFrac, ANOSIM *R* = 0.08, *P*_anosim_ = 0.014; Weighted UniFrac, ANOSIM *R* = 0.008, *P*_anosim_ = 0.016), but in-line with the observed high inter-individual variation within the groups, the low values of *R statistic* (<0.1) indicate that this was a small effect.

The within-group and overall high inter-individual variation observed in the beta diversity analysis (Fig. 1B-C) led us to do pairwise comparison of community dissimilarity between the control and ILI groups, as a means to control for high variability across all samples, and to potentially identify specific differences between groups. The microbiota of samples from the ILI group at the acute phase were significantly dissimilar to controls and recovery samples (Table 2). In addition, when compared to controls and recovery samples, samples from acute-ILI and ILI-14 days had significantly higher inter-individual variation in the NP microbiota (Wilcoxon rank-sum test, *p*.*adj* <□0.001, Fig. 1B). In general, in pairwise comparisons between groups, such as the control and ILI samples at the acute phase, the values of *R statistic* were higher than those obtained from comparisons within the ILI group, such as between recovery and acute samples (weighted Unifrac and GUnifrac). Overall, using different methods (see table 2) these data suggest that the changes in NP microbiota during acute-ILI can be highly variable between individuals, and potentially indicative of an unstable environment, which is greatly reduced after recovery.

**Table 2:** Pairwise comparisons of community dissimilarity between the groups. Analysis of similarity (ANOSIM) comparisons were based on weighted, generalized and unweighted UniFrac distances.

| Method | Group 1 | Group 2 | R | P-value |  |
| --- | --- | --- | --- | --- | --- |
| Weighted UniFrac | Control | Acute-ILI | 0.03682 | 0.001 | ** |
|  |  | ILI 14 days | 0.00437 | 0.196 |  |
|  |  | ILI-Recovery | 0.00177 | 0.284 |  |
|  | Acute-ILI | ILI 14 days | 0.00395 | 0.247 |  |
|  |  | ILI-Recovery | 0.01213 | 0.046 | * |
|  | ILI 14 days | ILI-Recovery | -0.00256 | 0.743 |  |
| GUniFrac | Control | Acute-ILI | 0.03113 | 0.001 | ** |
|  |  | ILI 14 days | 0.00624 | 0.136 |  |
|  |  | ILI-Recovery | 0.00303 | 0.211 |  |
|  | Acute-ILI | ILI 14 days | 0.00439 | 0.232 |  |
|  |  | ILI-Recovery | 0.01276 | 0.04 | * |
|  | ILI 14 days | ILI-Recovery | -0.00122 | 0.553 |  |
| Unweighted UniFrac | Control | Acute-ILI | 0.01081 | 0.024 | * |
|  |  | ILI 14 days | 0.01664 | 0.011 | * |
|  |  | ILI-Recovery | 0.00423 | 0.205 |  |
|  | Acute-ILI | ILI 14 days | 0.00857 | 0.116 |  |
|  |  | ILI-Recovery | 0.00604 | 0.129 |  |
|  | ILI 14 days | ILI-Recovery | 0.00141 | 0.278 |  |

### NP microbiota associations with ILI status

To identify compositional differences in microbial community members, we compared their relative abundances between the control and ILI groups. We observed high inter-individual variation in relative abundances of the dominant phyla, as seen in the distribution pattern for each of the groups (Fig.2A). At the phylum level, when compared to the control group, the microbiota of acute-ILI and 14-days after ILI had lower abundance of Actinobacteriota (Wilcoxon rank-sum test, *p*.*adj*□<□0.05, Fig. 2A). Proteobacteria were higher in samples from acute-ILI and ILI-14 day compared to controls, while samples from the recovery phase showed no significant differences when compared to controls.

**Figure 2:**
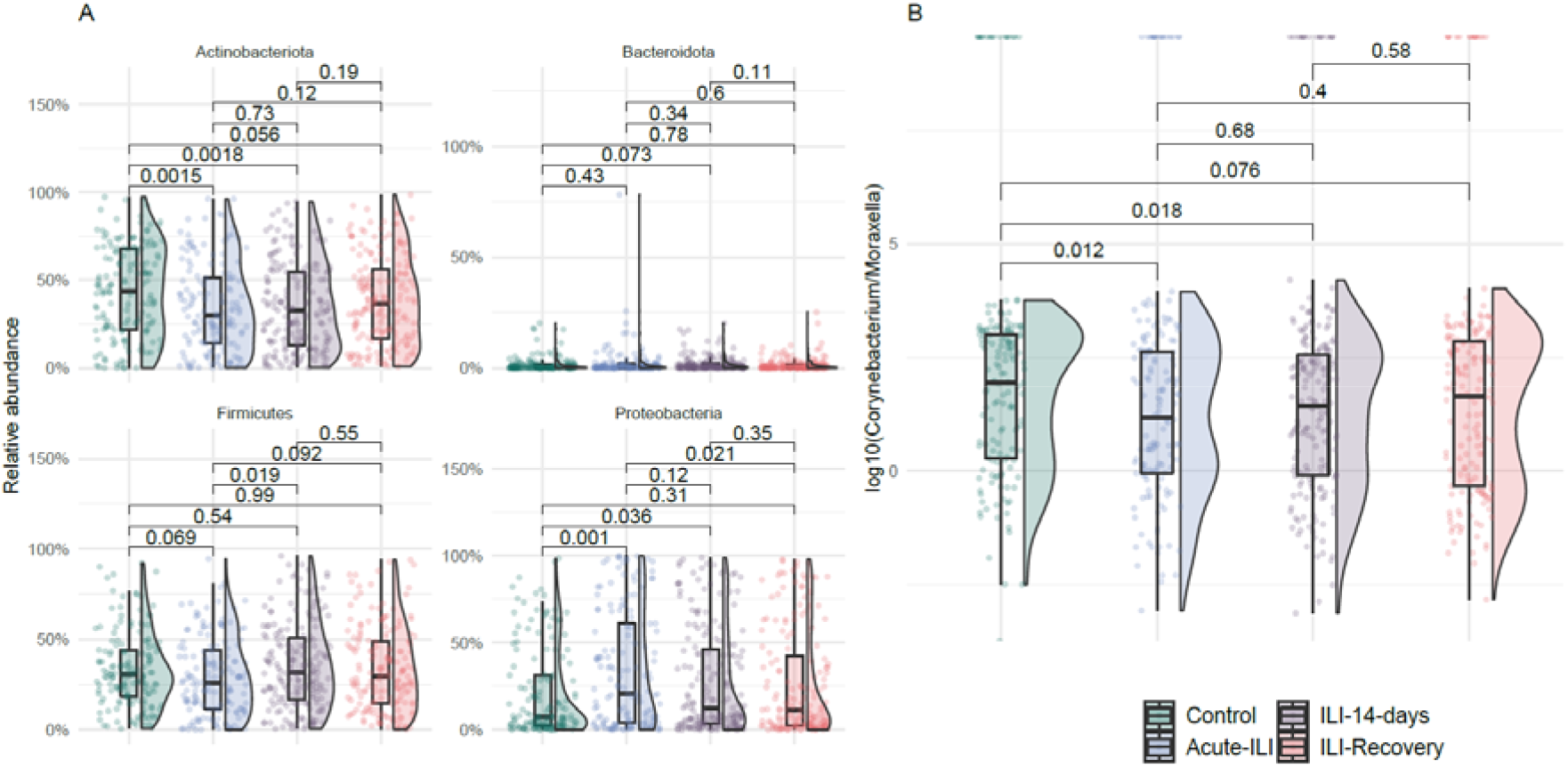
Comparison of phylum level abundances and *Corynebacterium*/*Moraxella* ratio. A] Relative abundances of the top four phyla were compared between groups using Wilcoxon rank-sum test, corrected for multiple comparisons using Benjamini-Hochberg Procedure. B] *Corynebacterium*/*Moraxella* relative abundance ratio was compared between groups using Wilcoxon rank-sum test, corrected for multiple comparisons using Benjamini-Hochberg Procedure.

Overall, the two most dominant genera, *Corynebacterium* and *Moraxella* showed a negative correlation (Spearman’s *rho*= -0.3728436, *P* = < 2.2e-16) potentially indicative of an anti-occurrence of these groups. Comparison of the *Corynebacterium*/*Moraxella* ratio suggested a significantly higher ratio in the control group when compared to acute-ILI and ILI-14 days but not with the ILI-recovery group (Fig. 2B).

We then tested for associations between the NP microbiota with having an ILI. Due to the intrinsic differences in our study groups, all associations were corrected for age and sex as potential confounders. Four genera were differentially abundant between the control samples and samples from acute-ILI. Among these, *Corynebacterium* showed a negative association, while *Haemophilus, Gemella* and *Porphyromonas* showed a positive association with acute-ILI (Fig. 3A and C). *Corynebacterium, Haemophilus*, and *Gemella* were not associated with other factors known to affect the upper respiratory tract microbiota, such as medication, smoking or other co-morbidities (including respiratory or cardiac disease(s), diabetes, kidney transplant, autoimmune diseases, asplenia, leukemia, lymphatic cancer and/or malignancies). However, *Porphyromonas*, positively associated with ILI, was negatively associated with consumption of alpha- & beta-blockers, other co-morbidities and respiratory diseases. Due to its association with medication and co-morbidities, its higher abundance in acute-ILI could not be directly associated with the acute-ILI event.

**Figure 3:**
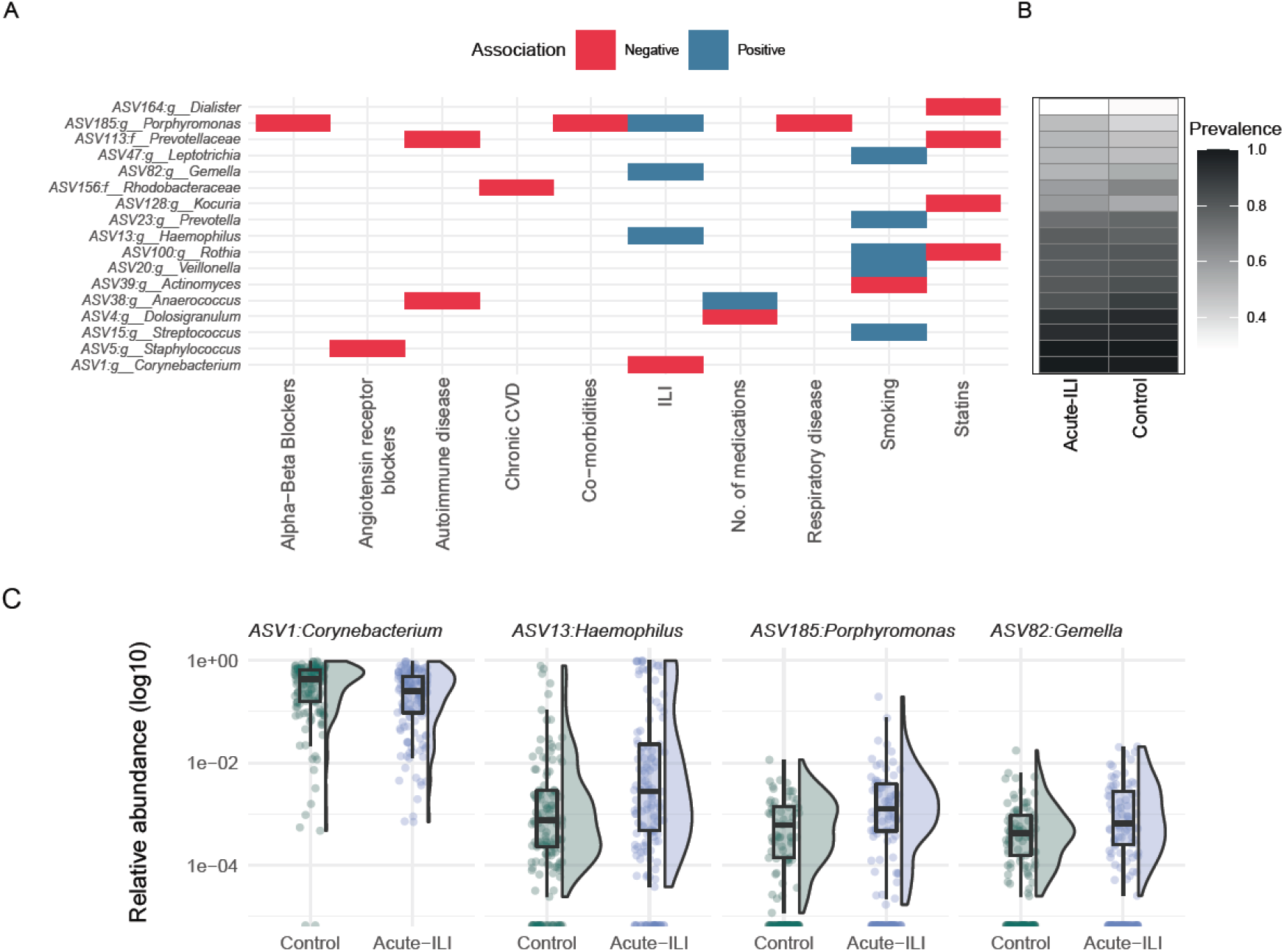
Genera associated with ILI status, medications and demographics. A] Genus-level associations with an FDR < 0.05. B] A heatmap showing prevalence of each of the genera in acute-ILI and control groups. C] Comparison of relative abundances of four genera associated with ILI status. *Haemophilus* infection (assessed by positive culture) was positively associated with *Haemophilus* relative abundances at the acute-ILI phase, with infected individuals showing high abundance even at the 14-day sample, and abundances after recovery were largely reduced (Supplementary Fig. 6).

### NP microbiota associations with health-related parameters independent of ILI

Our study population consists of older adults who are diagnosed with a variety of disease/disorders and are using medications that could potentially act as confounders. Therefore, we investigated potential associations with NP microbiota and other health related parameters and medications. *Staphylococcus* abundances were negatively associated with Angiotensin receptor blocker, while *Anaerococcus* and Prevotellaceae were negatively associated with autoimmune disease. Several genera found negatively associated with statins had low prevalence in both ILI (acute phase) and control groups (Fig. 3B). Smoking in the last 3 months was positively associated with relative abundances of *Streptococcus, Vellionella, Rothia, Prevotella* and *Leptotrichia*, however, the latter had low prevalence in both groups. These data suggest the need for investigating potential interactions between medications and NP microbiota, of especial interest in at-risk populations such as older adults, with high prevalence of co-morbidities and polypharmacy.

**Figure 3:**
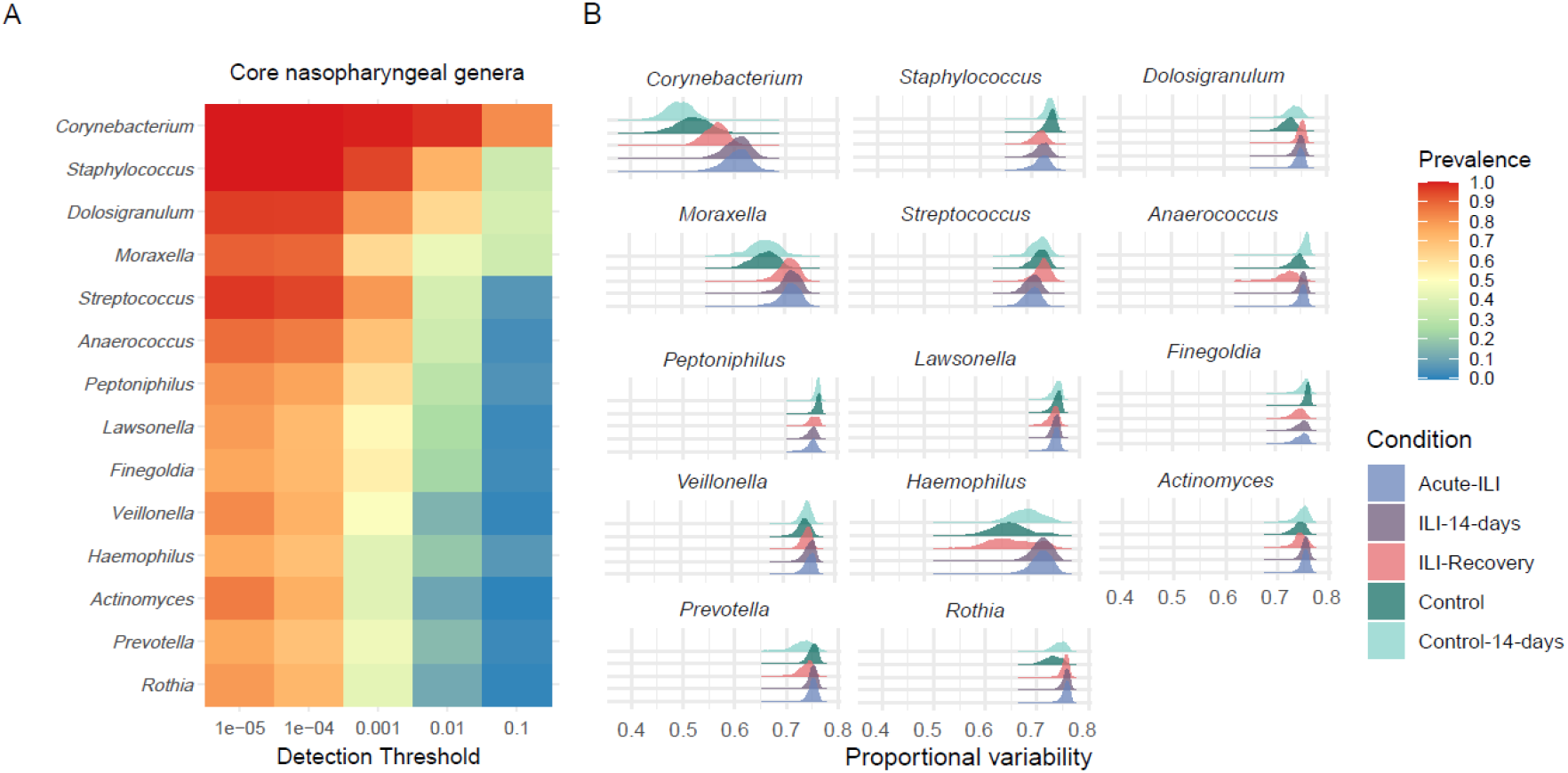
Core genera and their variation in different groups. A) Core genera identified in nasopharynx. B) Distribution of proportional variability values for each core genera within each group based on 999 bootstrap iterations.

#### Dynamics of the NP core microbiota during ILI

In host-associated microbial communities, members of the core microbiota can be viewed as stably-associated with the host. Since we observed high within-group variability in our study groups, we focused our analysis on the dynamics of the core-microbiota. We hypothesized that, in absence of ILI, the members of the core microbiota may exhibit lower inter-individual variability compared to acute-ILI, and can serve as a marker for recovery from the disease. For this analysis, we used paired samples of participants from the control (two visits, n=78) and ILI group (3 visits, n=81). This allowed us to investigate variability in core microbiota abundances between visits in each of the study groups. We identified 14 genera present in 75% of samples with at least 0.00001% relative abundance (Fig. 3A). Within core genera members such as *Anaerococcus, Peptoniphilus, Corynebacterium, Rothia* and *Prevotella*, variation in abundances at the ASV level was observed between the ILI (acute phase) and control groups (Supplementary Fig. 7). To test whether core genera showed variation in their abundance depending on disease status, we calculated the variation in the relative abundance of each core genera within a group using proportional variability (PV).. Comparison of proportional variability of core genera between groups revealed higher variation in abundances of the more dominant and prevalent genera, i.e., *Corynebacterium, Dolosigranulum, Haemophilus, Moraxella*, and *Rothia*, during of the acute phase of the ILI event (Fig. 3B).

Notably, the PV values for *Corynebacterium* and *Haemophilus* genera within the ILI samples at the recovery phase were more similar to the control group. On the contrary, *Staphylococcus* and *Streptococcus* had lower variability in ILI groups when compared to controls. These data suggest that the onset of ILI is characterized by variation in abundances of the dominant core genera, which is largely restored after recovery from the disease.

### Stability of the NP microbiota with and without ILI

Longitudinal stability of the microbiota is associated with resistance to pathogens and resilience to perturbations and is often suggested to have a positive association with diversity. We therefore investigated the relationship between stability of the NP microbiota with disease status and phylogenetic diversity. To this end, we calculated stability, defined as 1-GUniFrac dissimilarity, between paired samples in the absence of ILI (n=79) and during an ILI (n=81). No significant difference in stability was observed between visits for the control and ILI groups (Fig. 4A). However, in both groups there was a large inter-individual variability in microbiota stability between visits (Fig. 4 A and B). Notably, in the control group there was a negative correlation between phylogenetic diversity (PD) on the first visit and microbiota stability (Fig. 4C), also observed between samples from acute-ILI (v1) compared with 14 day and recovery visit microbiota.

**Figure 4:**
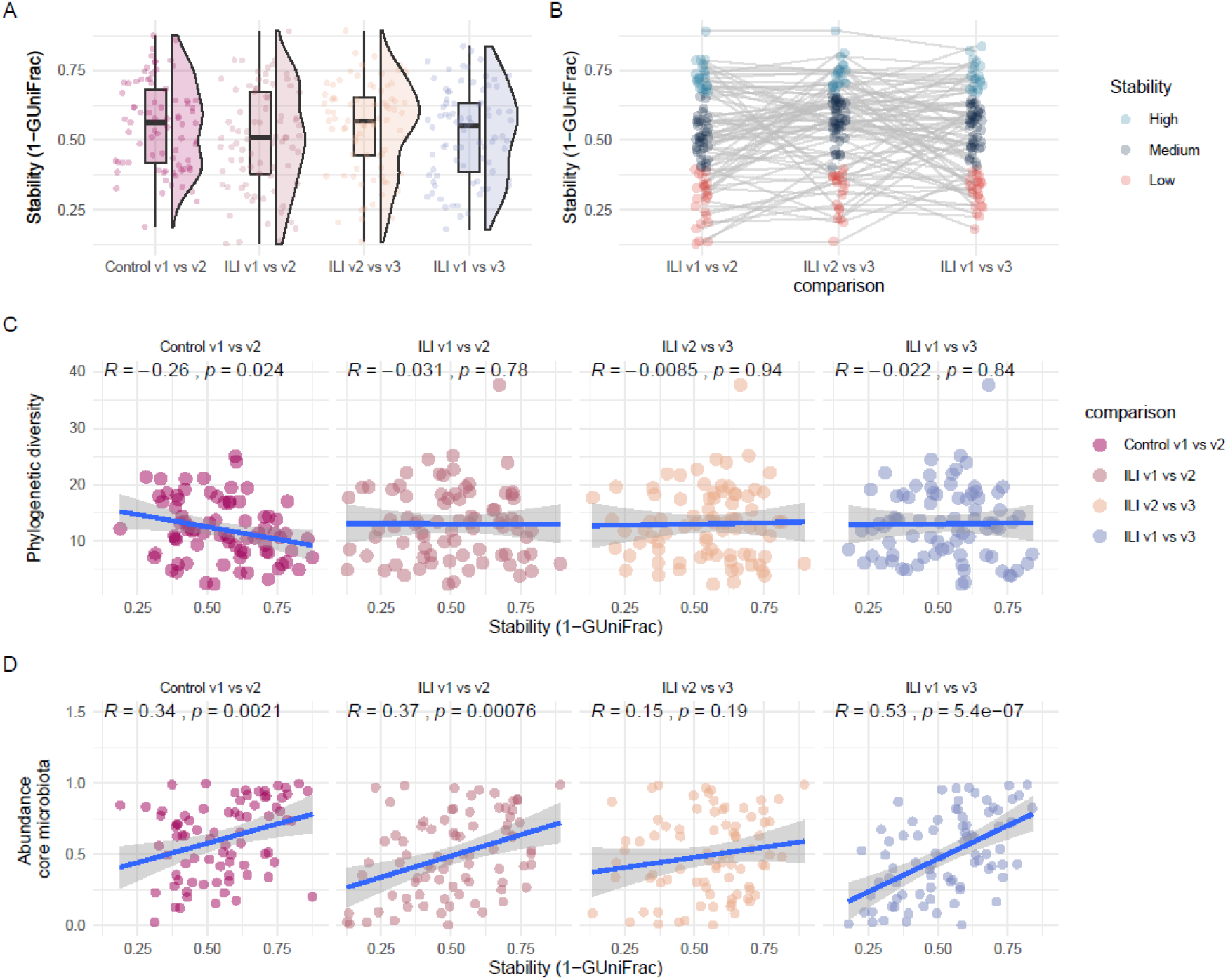
Associations of Stability and Diversity. A] Comparison of microbiota stability between visits. Here for control group, visits are randomly carried out throughout the influenza season, with 14 days intervals between first (v1) and second (V2) visit. For the ILI group, v1 is the acute-phase of ILI, v2 is 14 days later and v3 is visit between 7–9 weeks after v1. B] Depicts individuals categorized into three groups based on the stability of microbiota between visits. Each line connects observations for an individual to demonstrate differences between visits. C] Spearman’s correlation between stability (1-GUniFrac) and Phylogenetic diversity. D] Spearman’s correlation between stability (1-GUniFrac) and relative abundance of core microbiota.

Analysis of proportional variability indicated that the acute-ILI was associated with higher variability in relative abundances of key core microbiota members compared to controls. We further investigated whether the observed microbiota stability was associated with the relative contribution of core members (core abundance) to the total community abundance. We observed that higher core abundance was associated with higher stability between visits in the control group, as well as between acute-ILI (visit-1) with 14 days (visit-2) and with recovery phase (visit-3) respectively (Fig. 4D). However, core abundances did not correlate with stability between 14 days (visit-2) and recovery stage (visit-3), indicative of a more unstable environment in the early phase of recovery from the disease.

## Discussion

Here we studied the NP microbiota in older adults during an influenza like illness event. We evaluated the differences in the NP microbiota in those not reporting an ILI event throughout the 2014-2015 influenza season with those reporting an ILI event, sampled during the acute phase of ILI (<72h after report), 14 days after and 7–9 weeks after the onset of ILI, considered as the recovery phase. Acute ILI was associated with differences in microbiota community and structure when compared to controls and recovery samples. The NP microbiota exhibited high inter-individual differences in dynamics from onset to recovery from ILI. Dominant core genera showed higher variability in acute-ILI and 14 days after ILI when compared to controls, indicative of an unstable microbiota environment in the early days of reporting an ILI event.

The low-biomass nature of NP swabs used in this study presents a major challenge in separating exogenous taxa from “true” observations in the samples. Previously, several researchers have highlighted this issue and proposed potential approaches to tackling the “kitome” in low-biomass microbiome research [32-35]. Iteratively going through every step, i.e., decontamination using decontam package, checking for abundances in negative controls, and identifying prevalent and dominant contaminants that were abundant in both controls and samples using co-occurrence analysis, aided in removal of a large number of potentially contaminating taxa. While the decontam package identified several contaminants, requirement of additional steps after decontamination have been recently reported, and in our study, we also noticed the importance of detailed investigation of contaminants [27, 49]. Furthermore, taxonomic investigation as well as looking for isolation sources for doubtful taxa in well curated public resources like the Bacterial Diversity Metadatabase (BacDive) proved as a useful tool for rational choice on filtering [31]. We sequenced 45 negative controls (32 from the DNA extraction step and 13 from the PCR step) which allowed us to identify the DNA extraction step as the main source of introduction of contamination [34]. Resources such as BacDive, and ecosystem specific databases such as the Human Oral Microbiome Database, are useful for improving confidence in observed taxa [31, 50]. Our observations with NP swabs highlight the need for including both positive and negative controls at every step possible for rigorous quality control, and a downstream analysis that is driven by skepticism of detected taxa.

Micro-niches within the upper-respiratory tract are occupied by distinct microbial communities and diverse bacteria can co-exists in the NP [8, 9]. Anatomically, the NP is located above the oral cavity and can be blocked during respiratory infections due to nasal congestion. Moreover, local changes in inflammatory responses may influence the habitability of NP by microbes. Thus, changes in these micro-niches can occur during respiratory infections thereby potentially impacting the commensal microbiota through proinflammatory responses or shortage of nutrients [51, 52]. Acute-ILI was characterized by higher abundances of Proteobacteria and lower abundances of Actinobacteriota and Firmicutes. Similar observations have been previously reported in individuals with upper respiratory tract viral infections [15]. Notably, Actinobacteriota and Firmicutes had higher abundances after recovery from ILI, further suggesting that these phyla are negatively associated with ILI at the acute phase and are largely restored after disease [53]. At the genus level, the *Haemophilus, Gemella* and *Porphyromonas* genera were positively associated with acute-ILI. *Haemophilus* was more abundant in infected individuals compared to non-infected individuals within the ILI group at the acute phase. *Haemophilus* was also more abundant in samples collected 14 days after the acute-ILI and only reduced after recovery from the ILI, likely suggesting a reduction of the pathogenic variant at a later phase of recovery from ILI. Previously, *Haemophilus influenzae* was detected in samples collected at the recovery phase as well as in control samples in this cohort [6]. Species or strain typing for *H. influenzae* or *H. haemolyticus* is challenging using short-amplicon sequencing approaches as used in this study, however, the overall positive association confirms the biological signal captured in our analysis. The genus *Gemella* is widely considered as a commensal of the respiratory niche, however, there is evidence for it being a potential opportunistic pathogen in older adults [54]. Similarly the genus *Porphyromonas*, more abundant in acute-ILI compared to controls, has been associated with pro-inflammatory responses [55, 56]. Notably, the gut microbiota of individuals with acute-ILI was previously reported to harbor a pro-inflammatory profile [17]. Overall, based on our observations we hypothesize that ILI might promote expansion of pro-inflammatory bacteria, as previously shown for the gut microbiota, also locally within the respiratory tract. In addition, previous studies have observed an association between the nasal microbiota and inflammatory responses during infections [9, 57]. Therefore, future studies should investigate the dynamics interactions between immune system and microbiome associated with ILI.

*Corynebacterium*, that belongs to the Actinobacteriota phylum, was the only genera that had a negative association with acute-ILI. *Corynebacterium* is proposed as one of the keystone species in the upper respiratory tract, mainly due to its positive association with health [8, 58]. The majority of the previous studies of the NP microbiota included children and adults (<65 years age), but a recent study during influenza virus infection across age-groups (including older adults, n=66) observed a lower prevalence of *Corynebacterium* [16]. Our observation of lower abundances *Corynebacterium* during ILI at the acute phase, and its increase after recovery from the disease in participants >60 years of age, indicates a potential life-long key role for *Corynebacterium* and related species with health in the upper respiratory tract. Future studies investigating the mechanistic role of *Corynebacterium* in the upper respiratory tract are need to better understand the impact of this bacterium in human health. In a previous study, it was suggested that core upper respiratory tract microbiota is perturbed during influenza A virus infection [15]. The core microbiota represents taxa that are highly prevalent, likely due to their adaptation to a particular ecosystem, and could play a vital role in maintaining stability and therefore health [59, 60]. Hence, any variation in the core microbiota could suggest imbalances in the ecosystem, leading to susceptibility for disease or simply serving as a biomarker of disease. Therefore, we compared the variability in the core microbiota in our cohort. To this end, we quantified the inter-group variability in core microbiota using a non-parametric measure, Proportional Variability (PV) [41, 42]. PV can be a better estimate of variability for quantities undergoing very different dynamics, in our study, relative abundances of core bacterial taxa. Although it is difficult to estimate a range of abundance values for bacteria that can be considered as “healthy”, variability within groups can serve as markers for perturbation, as suggested in the analogy with Anna Karenina principle [61]. We observed higher variability within the ILI group at the early phases (acute and 14 days) for dominant and prevalent taxa such as *Corynebacterium, Moraxella* and *Dolosigranulum*, which have been previously associated with a “healthy” NP microbiota [8]. Lower variability of core genera such as *Streptococcus* and *Staphylococcus* was observed in the ILI group compared to the control group. This indicates that some core bacteria are strongly or similarly selected during ILI in the majority (if not all) of the diseased participants, thus showing lower variability within the diseased group. Furthermore, pathogenic strains within *Streptococcus* and *Staphylococcus* genera can cause complications and exacerbate respiratory infection [62]. There is a need for investigating strain-level variation between pathogenic (non-core) strains and core strains of *Streptococcus* and *Staphylococcus*, which could illuminate the potential impact of strain variation on host-susceptibility to infection.

The longitudinal analysis of beta-diversity indicated that the stability (1-GUniFrac) of the overall microbial community was variable between visits, independently of the disease status. A negative association between phylogenetic diversity, a measure of biodiversity, and stability in the absence of ILI suggests that individuals with higher phylogenetic diversity exhibit a more variable microbiota composition. This is contrary to the widely reported diversity -stability relationship and requires further investigation in the context of upper respiratory microbiota and its importance in host-health. The abundance of core microbiota was associated with higher stability in controls and also between acute-ILI and other visits. This suggests that the core microbiota at the acute phase of ILI potentially plays a critical role in stability and therefore recovery of the NP microbiota, especially 7-9 weeks after the onset of ILI.

## Conclusion

Investigation of the NP microbiota in the older adult population suggests that onset of ILI is accompanied by changes in the microbiota. Potential pro-inflammatory bacteria increase in abundances during acute-ILI. Some strains belonging to genera such as *Haemophilus, Streptococcus* and *Staphylococcus* consists of strains that are known causative agents of secondary infections related to complications of ILI in the ageing population [63]. Increased abundances of *Gemella* and *Prophyromonas* in acute-ILI, suggests that there is a need to investigate the role of these bacteria during ILI. Variability in core microbiota can be viewed as a potential biomarker of ILI, indicating that higher core microbiota abundance during onset of ILI is associated with higher resilience to changes during recovery stage. Future studies should focus on the role of core NP microbiota in determining the susceptibility to ILI and its influence on recovery. For this, a trans-domain integrated approach will be crucial to better understand the three-way interaction between core microbiota-host immune system-viral infection outcome.

## Availability of data and material

Raw sequencing data analyzed in this study are deposited at European Nucleotide Archive database under the study accession number PRJEB46215. The R code and scripts used to analyze the data are available from the GitHub repository https://github.com/RIVM-IIV-Microbiome/ILI-Respiratory-Microbiota-2021. Information pertaining to participant data is available following institutional regulations.

## List of abbreviations

NP: Nasopharynx
ILI: Influenza-like illness
rRNA: Ribosomal ribonucleic acid
OP: Oropharynx
PCR: Polymerase chain reaction
qPCR: Quantitative Polymerase chain reaction
ATCC: American Type Culture Collection
ASV: Amplicon sequence variant
NMF: Non-negative Matrix Factorization
BacDive: The Bacterial Diversity Metadatabase
GUniFrac: Generalized UniFrac
PV: Proportional variability
COPD: Chronic obstructive pulmonary disease
PD: Phylogenetic diversity
ANOSIM: Analysis of similarities

## Acknowledgements

This work was supported by the Dutch Ministry of Health, Welfare and Sport, and the Strategic Program of the National Institute for Public Health and the Environment (RIVM). We thank all participants for their invaluable contribution to this study.

## Author Contributions

DvB and JvB were responsible for the ILI study. DvB, and SF conceived and designed respiratory microbiota experiments. TB coordinated and JS and SK performed laboratory work. SAS performed the microbiota analysis. SAS wrote the initial draft of the manuscript with inputs from SF. All authors contributed to interpreting the results, critically revisions of the manuscript and approved the final manuscript.

## Ethics declarations

### Ethics approval and consent to participate

Ethical clearance was obtained from the Medical Ethical Committee Noord Holland and written informed consent was provided by the participants. This study is registered with the Netherlands Trial Registry, number NL4666.

### Competing interests

The authors have no competing interests.

## Notes

### Competing Interest Statement

The authors have declared no competing interest.

