## Supplementary information for "Associations and recovery dynamics of the nasopharyngeal microbiota during influenza-like illness in the aging population"

**Supplementary text:**

### Dealing with contaminant sequences

Investigation of taxonomic composition of technical controls indicated a high prevalence of taxa previously reported as reagent contaminants, e.g. *Ralstonia*, *Bradyrhizobium*, *Mesorhizobium*, *Comamonadaceae* (Supplementary fig. 1). These contaminant taxa also dominated several NP samples, suggesting the need for strict evaluation and identification of contaminants prior to our analyses (Supplementary fig. 2). As a first step to remove contaminating ASVs, we used the prevalence methods in the decontam R package function *isNotContaminant()* using a 0.5 threshold. These resulted in removal of 4.8% of the total reads in the data, contributed by 5706 ASVs. After this, further investigation of ASVs indicated presence of several known potential contaminants [1-4]. Therefore, we included a second decontamination step, where we removed ASVs over-abundant in negative controls compared to controls, such as those classified as Cyanobacteria, Chloroflexi and Mitochondria, as well as samples with less than 2000 reads. At this step 474 ASVs were removed which comprised 2.5% of the total reads.

Next, we evaluated ASVs that were identified in negative controls and subset these from all the samples (referred to as blank-sample ASVs). We noticed that ASVs classified as *Corynebacterium*, *Moraxella*, *Dolosigranulum*, *Streprococcus*, known inhabitants of upper respiratory tract, were present in some negative controls. These were mostly dominating the DNA extraction control and likely a result of cross-contamination from NP samples during sample processing. Using a co-abundance approach based on Spearman's correlation, we identified 9 co-abundance clusters. Randomly splitting the entire data found good agreement between the created test and training sets (r: 0.769 and significance: 0.001). Investigation of taxonomic identities of the ASVs in each of the clusters, highlighted taxa that are highly prevalent and abundant in negative controls dominated cluster 4, 7, 8 and 9 (Supplementary table 2). The cluster 4 was dominated by ASV7:*Burkholderia-Caballeronia-Paraburkholderia*, ASV8:*Comamonadaceae* and ASV9:*Burkholderiales* ASVs. The cluster 7 was dominated by ASVs classified as *Methylobacterium-Methylorubrum*, cluster 8 and 9 were dominated by low abundant and prevalent ASVs mostly known to be potential contaminants. All the ASVs belonging to cluster 4, 7, 8 and 9 were removed from the dataset for further analysis. Overall, during the decontamination process, ~20% of the raw reads were removed, and the final data used for microbiota analysis consisted of 3152 ASVs in 758 samples. These data were considered to have none (or minimal) contaminant ASVs to distort the proportional abundances or diversity-based comparisons, and association analysis.

**Supplementary figures**

**Supplementary Figure 1: Overview of the samples and analysis done in this study.**

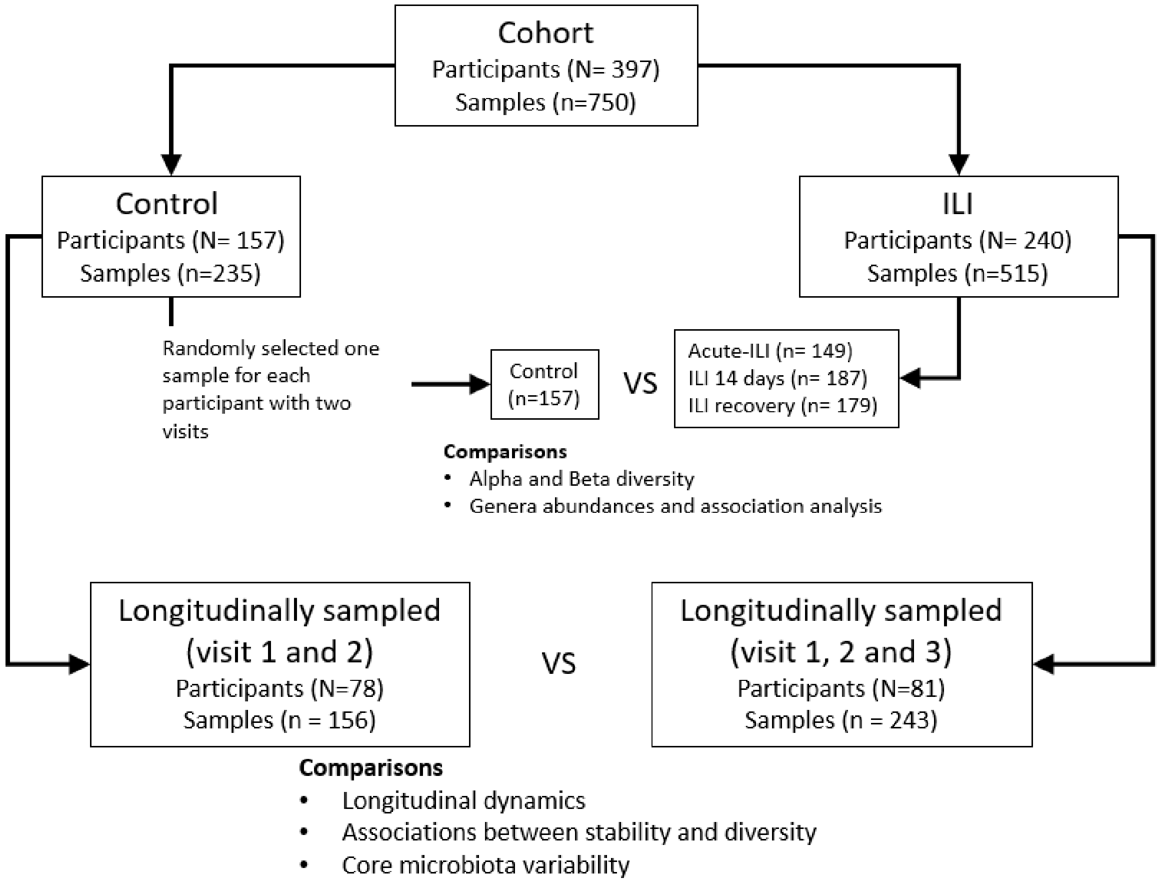

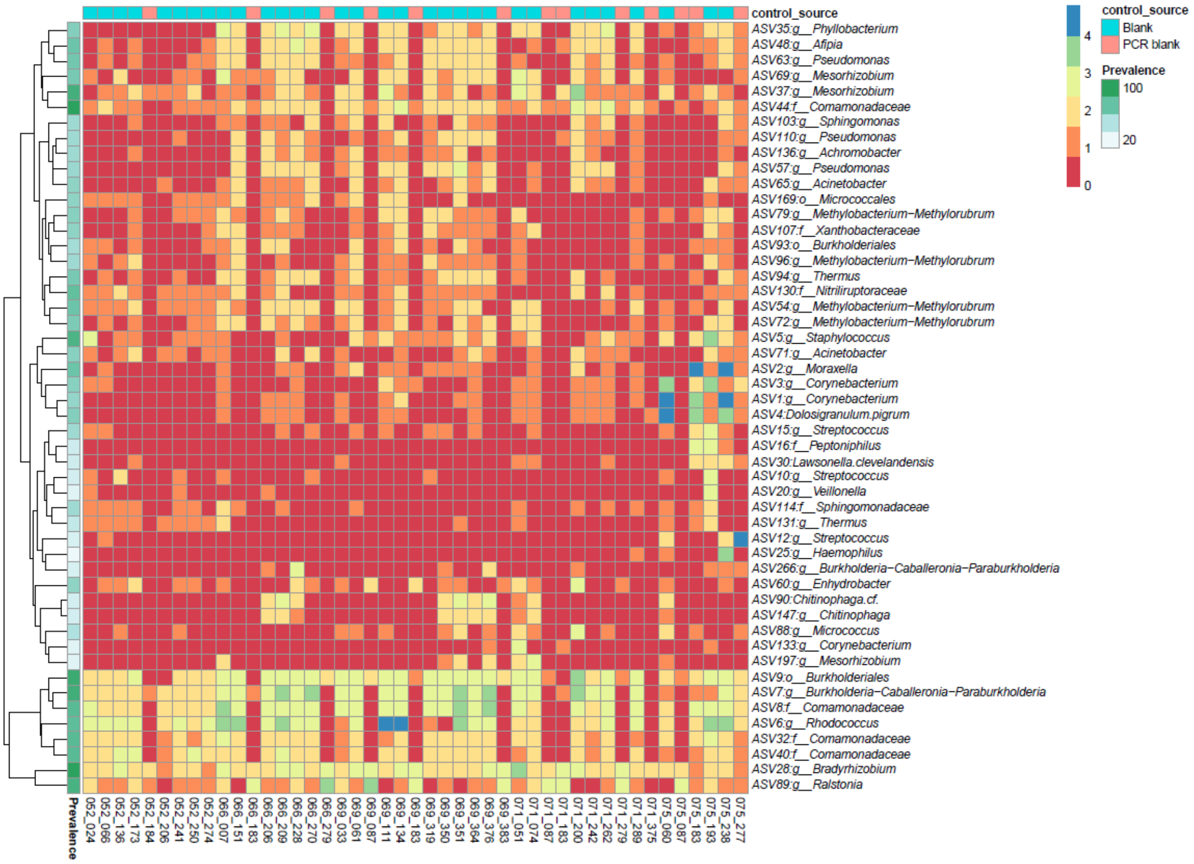


**Supplementary figure 2: Composition top 50 ASVs in negative controls.** Blank refers to DNA extraction negative control and PCR blank refers negative control for polymerase chain reaction. The color intensity is log10 +1 scaled for visualization.

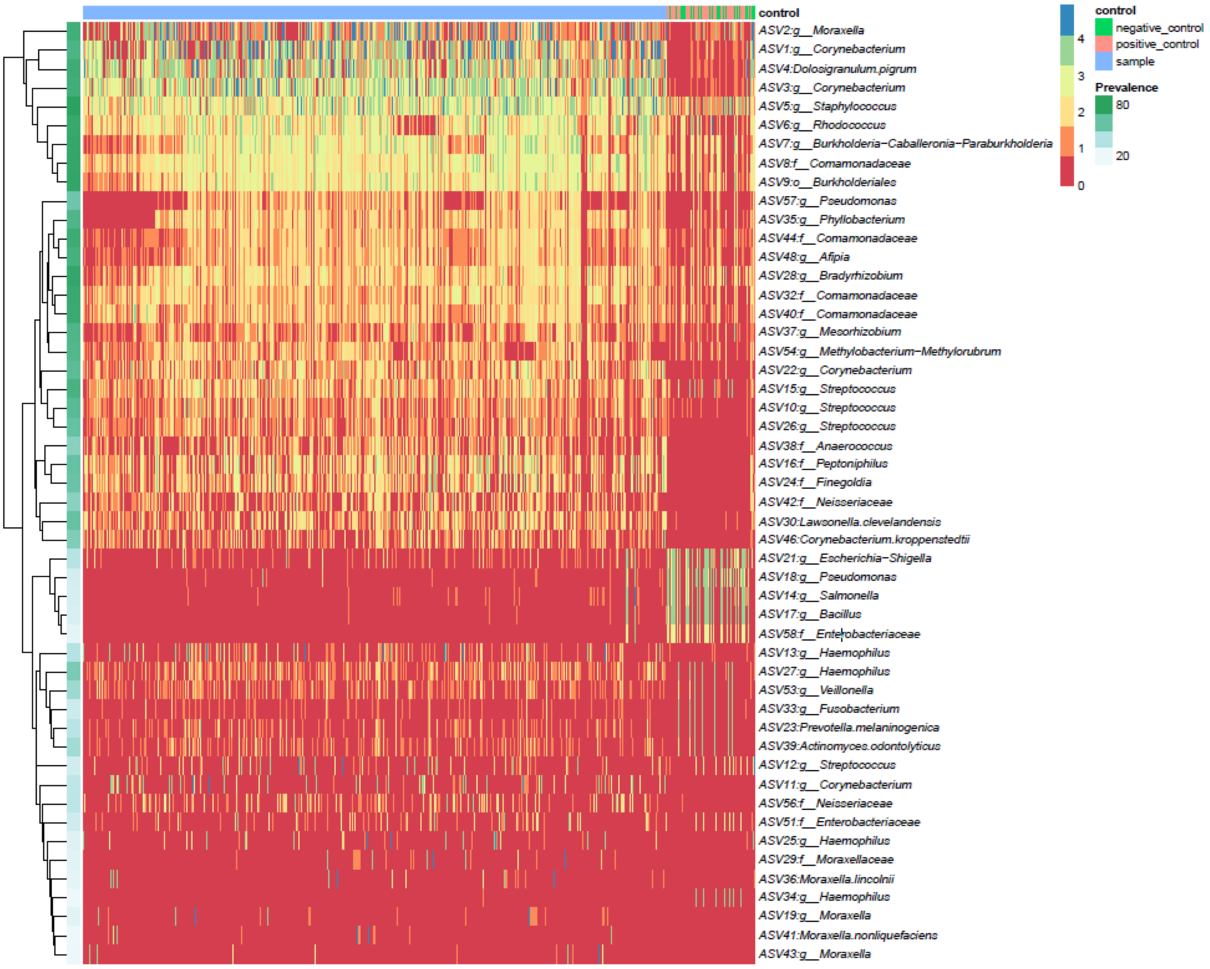


**Supplementary figure 3: Composition top 50 ASVs in technical controls and nasopharyngeal samples.** The color intensity is log10 +1 scaled for visualization.

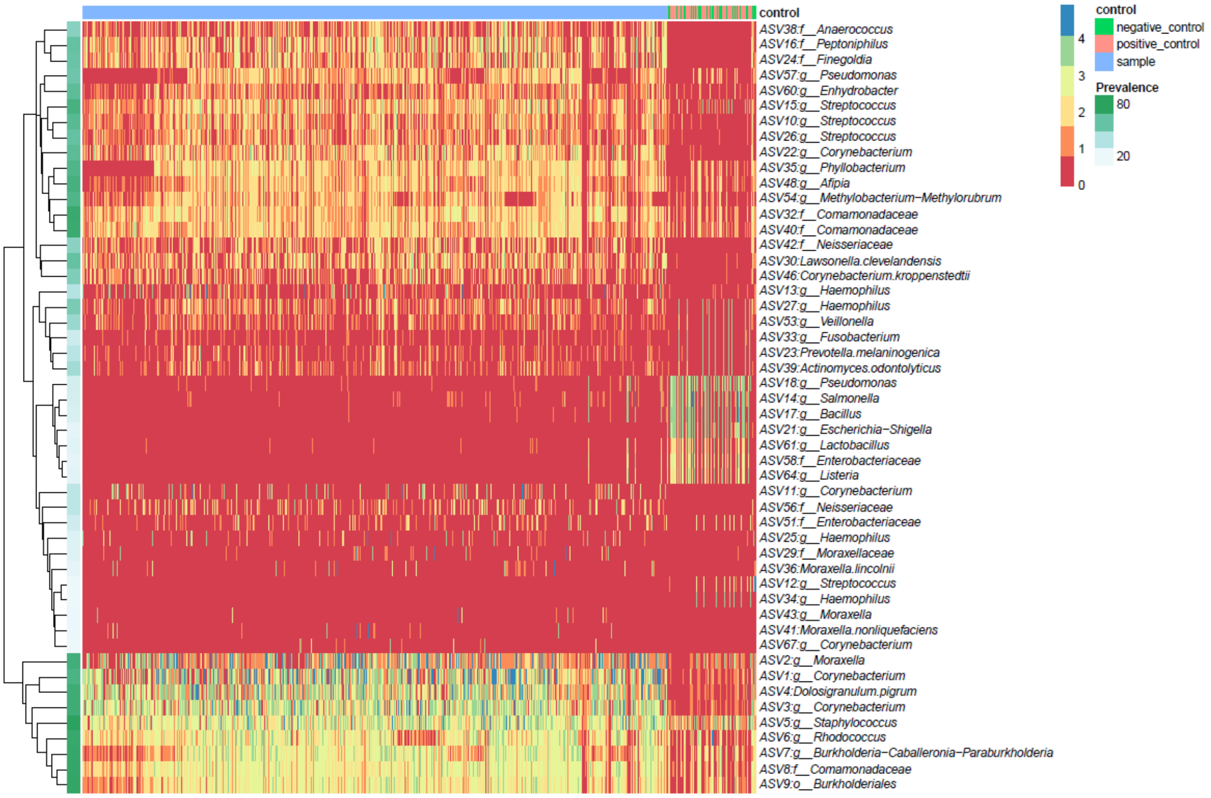


**Supplementary figure 4: Composition top 50 ASVs in technical controls and nasopharyngeal samples after removal by decontam's *isNotContaminant* function**. The color intensity is log10 +1 scaled for visualization.

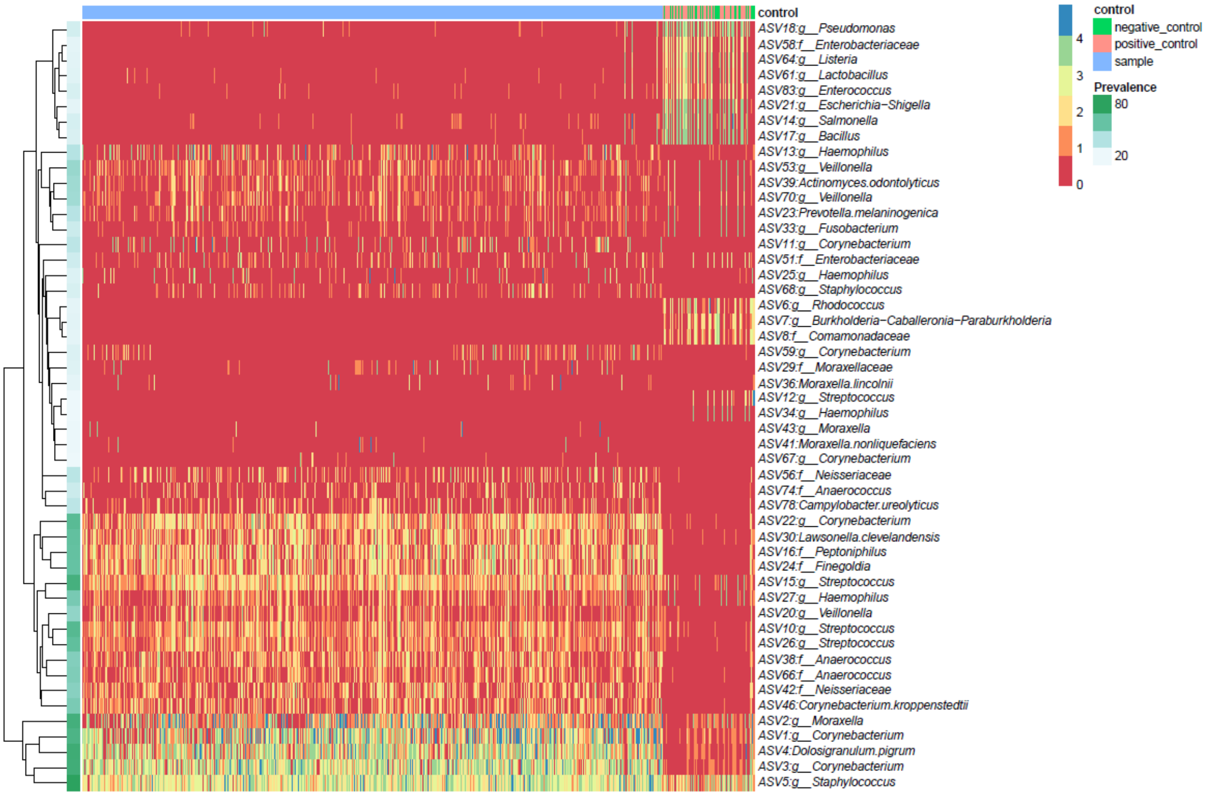


**Supplementary figure 5: Composition top 50 ASVs in technical controls and nasopharyngeal samples after removal of co-occurring clusters of contaminants**. The color intensity is log10 +1 scaled for visualization.

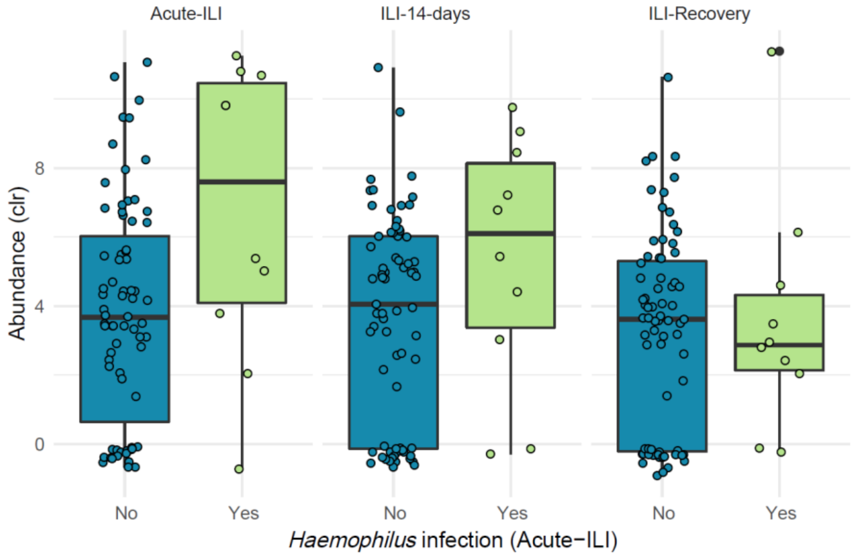


**Supplementary figure 6: Abundance of genus *Haemophilus* in ILI participants at different stages.** The infection status at acute-ILI stage is labelled on x-axis. Infection at acute-ILI=yes; no infection at acute-ILI=no. The abundance values are centered log-ratio (clr) transformed. Samples shown are those where we have information for all 3 sampling points.

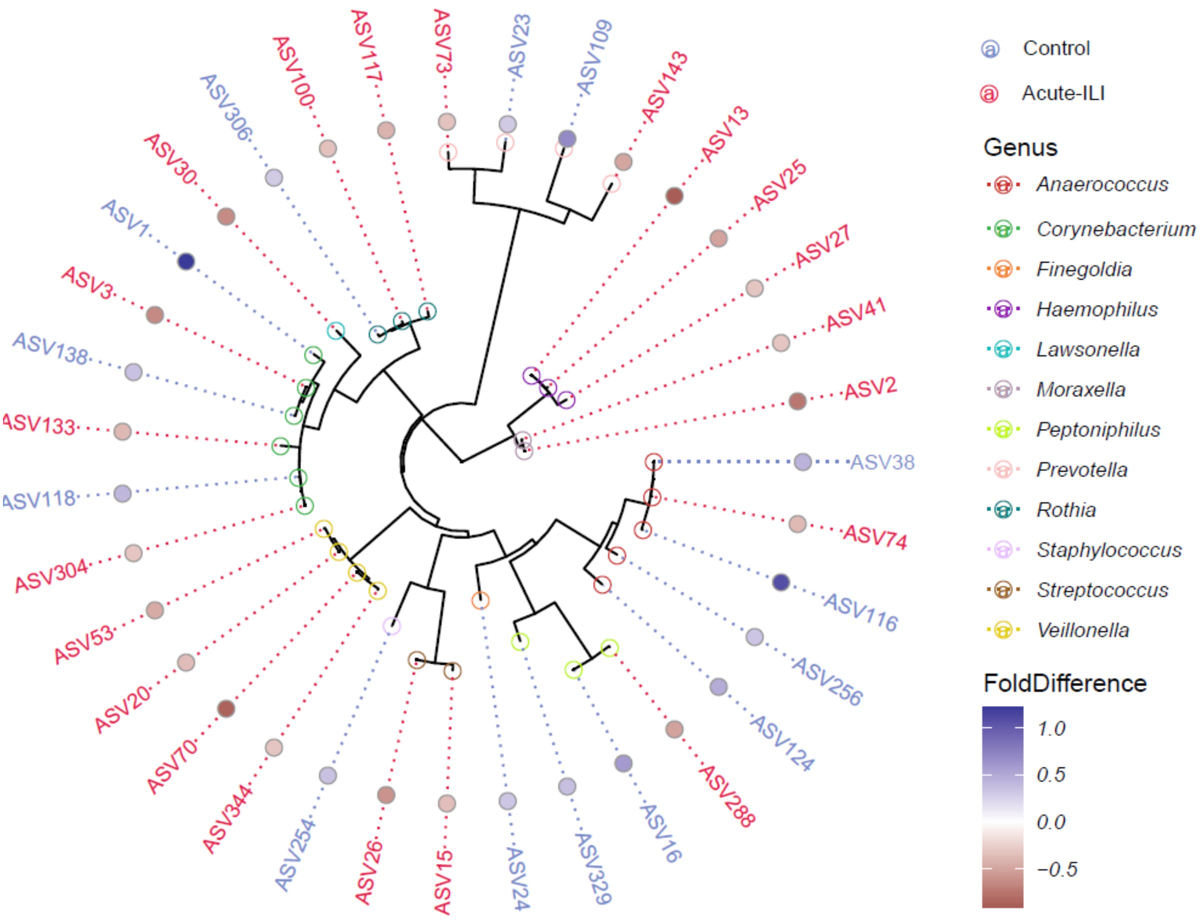


**Supplementary Figure 7: ASV level differences in core genera between control and acute-ILI.** A) ASVs part of the core genera were extracted and fold-difference. ASV labels are colored depending on their enrichment in controls or acute-ILI.

**Supplementary Table 1:** Pathogens detected by Multiplex ligation-dependent probe amplification (MLPA) or classical culture approaches in samples from ILI participants.

| **Pathogens** | **Method** | **ILI (N=238)** |
|---|---|---|
| Influenza virus | | 42/198 |
| Influenza A virus | MLPA | 25/215 |
| Influenza B virus | MLPA | 17/223 |
| Rhinovirus | MLPA | 48/198 |
| Coronavirus | MLPA | 19/221 |
| RSV | MLPA | 12/228 |
| HPIV | MLPA | 12/228 |
| HMPV | MLPA | 11/229 |
| Bocavirus | MLPA | 2/238 |
| Adenovirus | MLPA | 3/237 |
| *Mycoplasma pneumoniae* | MLPA | 8/232 |
| *Chlamydophila pneumoniae* | MLPA | 1/239 |
| *Haemophilus* | | 37/203 |
| ~*H. influenzae* | culture | 27/213 |
| ~*H. haemolyticus* | culture | 10/230 |
| *Haemolytic Streptococcus* | culture | 9/231 |
| *Staphylococcus aureus* | culture | 13/227 |
| *Moraxella catarrhalis* | culture | 1/239 |

**Supplementary table 3: Overview of phyla detected by 16S rRNA gene profiling.**

| Phylum | Counts | Percent | Prevalence (%) | Note | Total Percent |
|---|---|---|---|---|---|
| Actinobacteriota | 7254136 | 40.058 | 99.868 | Kept | 99.782 |
| Firmicutes | 5630742 | 31.093 | 100.000 | Kept | |
| Proteobacteria | 4834685 | 26.698 | 100.000 | Kept | |
| Bacteroidota | 263533 | 1.455 | 96.570 | Kept | |
| Fusobacteriota | 51169 | 0.283 | 66.095 | Kept | |
| Campilobacterota | 35294 | 0.195 | 48.813 | Kept | |
| Deinococcota | 27300 | 0.151 | 63.193 | Discarded | 0.218 |
| Synergistota | 2846 | 0.016 | 5.805 | Discarded | |
| Planctomycetota | 2164 | 0.012 | 20.185 | Discarded | |
| Acidobacteriota | 1736 | 0.010 | 20.844 | Discarded | |
| Spirochaetota | 1125 | 0.006 | 11.346 | Discarded | |
| Bdellovibrionota | 844 | 0.005 | 11.741 | Discarded | |
| Verrucomicrobiota | 825 | 0.005 | 8.707 | Discarded | |
| Gemmatimonadota | 662 | 0.004 | 10.026 | Discarded | |
| Desulfobacterota | 526 | 0.003 | 8.443 | Discarded | |
| Myxococcota | 498 | 0.003 | 3.958 | Discarded | |
| Crenarchaeota | 310 | 0.002 | 2.507 | Discarded | |
| Abditibacteriota | 279 | 0.002 | 3.430 | Discarded | |
| Euryarchaeota | 156 | 0.001 | 2.243 | Discarded | |
| Halobacterota | 114 | 0.001 | 0.792 | Discarded | |
| Armatimonadota | 103 | 0.001 | 1.583 | Discarded | |
| Fibrobacterota | 21 | 0.000 | 0.660 | Discarded | |
| Nitrospirota | 17 | 0.000 | 0.396 | Discarded | |
| Dependentiae | 8 | 0.000 | 0.264 | Discarded | |

**Supplementary table 4: Overview of top 10 genera detected by 16S rRNA gene profiling.**

| Genus | Counts | Percent |
|---|---|---|
| *Corynebacterium* | 6857479 | 37.95045 |
| *Moraxella* | 3468908 | 19.19752 |
| *Dolosigranulum* | 2263348 | 12.52575 |
| *Staphylococcus* | 1987030 | 10.99656 |
| *Haemophilus* | 574871 | 3.181433 |
| *Streptococcus* | 318384 | 1.761991 |
| *Peptoniphilus* | 273254 | 1.512234 |
| *Anaerococcus* | 252891 | 1.399542 |
| *Finegoldia* | 176282 | 0.975574 |
| *Prevotella* | 129608 | 0.717273 |
| **Top 10 Classified** | 16302055 | 90.21833 |
| **Unclassified** | 563054 | 3.116036 |

**Supplementary table 5: Dominant genera identified in the study.**

| Genus | No. Samples Genus dominates | Percent of total samples (%) |
|---|---|---|
| *Corynebacterium* | 375 | 49.5 |
| *Moraxella* | 154 | 20.3 |
| *Staphylococcus* | 104 | 13.7 |
| *Dolosigranulum* | 39 | 5.1 |
| *Haemophilus* | 30 | 4 |
| *Streptococcus* | 12 | 1.6 |
| *Salmonella* | 4 | 0.5 |
| *Anaerococcus* | 3 | 0.4 |
| *Brochothrix* | 3 | 0.4 |
| *Peptoniphilus* | 3 | 0.4 |
| *Prevotella* | 3 | 0.4 |
| *Neisseria* | 2 | 0.3 |
| *Finegoldia* | 1 | 0.1 |
| *Fusobacterium* | 1 | 0.1 |
| *Kocuria* | 1 | 0.1 |
| *Lactobacillus* | 1 | 0.1 |
| *Mycoplasma* | 1 | 0.1 |
